# The indoleamine 2,3 dioxygenase pathway drives intratumoral B cell maintenance

**DOI:** 10.1101/2021.08.25.456776

**Authors:** Burles A. Johnson, Adam K. Aragaki, Donna M. Williams, Ophelia Rogers, Jack Mountain, Li Luo, Wenhao Zhang, Lingling Xian, Mingxiao Feng, Lionel Chia, Dominic Dordai, Noah M. Hahn, Stephen Desiderio, Theodore S. Johnson, David J. McConkey, Linda M.S. Resar

## Abstract

B cells have been implicated as central regulators of immune responses in settings as diverse as mammalian pregnancy, mucosal tolerance, chronic infection states, autoimmunity, and the tumor microenvironment. Despite the established importance of B cells in these environments, the mechanisms by which B cells are maintained in these contexts remain undefined. Here, we report that IDO1 pathway inhibition with D-1-methyl-tryptophan (D-1MT) and linrodostat significantly decreases tumor infiltrating B (TIL-B) cells in a preclinical model of melanoma. Single cell RNA sequencing (scRNAseq) of murine melanoma demonstrate TIL-B cells are heterogeneous but primarily express markers consistent with an immune stimulatory phenotype. D-1MT decreases splenic B cells and bone marrow derived B cell precursors in tumor-bearing mice, suggesting that IDO1 pathway inhibition impedes B cell maturation. D-1MT decreases intratumoral myeloid derived suppressor cells (MDSCs), which are essential for maintenance of TIL-B cells. Unlike D-1MT, genetic deletion of tumor *Ido1* does not impact TIL-B or MDSC numbers. In human solid tumors, intratumoral *IDO1* expression consistently associates with high expression of a pan-B cell gene signature, and in patients with melanoma, scRNAseq analysis of tumor samples revealed most TIL-B cells express *IDO1*. Collectively, our data reveal the impact of pharmacologic IDO1 inhibition on B cells, which may have therapeutic implications for patients with solid tumors by informing the design of future oncology clinical trials.

## Introduction

B cells are a critical regulator of the immune response in settings such as pregnancy^1^, infection^2^, autoimmunity^3^, and cancer^4^. As B cells have diverse function such as antigen presentation, T cell activation, immunoglobulin generation, and cytokine production, these cells can exude powerful context dependent immune stimulatory or immune inhibitory effects^1–4^. In cancer, B cells can stimulate T cell responses to enhance anti-tumor immunity^5^; however, immune-regulatory B cells (Bregs) suppress responses^6,7^ and correlate with advanced tumor stage, disease progression, and/or decreased survival in diverse solid tumors^8–13^. An understanding of mechanisms governing intratumoral B cell maintenance may provide additional therapeutic targets to enhance stimulatory B cell function, and/or mitigate suppressive Breg effects.

Accumulating evidence suggests amino acid metabolism is critical for B cell function and survival in certain conditions^14,15^. While much focus has centered on glutamine metabolism in regulating B cell function^16^, other amino acid pathways can modulate B cells. For example, B cell specific expression of the tryptophan catabolizing enzyme indoleamine 2,3 dioxygenase-1 (IDO1) suppresses B cell proliferation and antibody secretion in response to T cell-independent antigen stimulation^17^. Conversely, IDO1 activity facilitates autoantibody production in a model of gastric dysplasia^18^. Thus, the role of IDO1 in B cells is likely complex and context dependent. B cells play an important role in regulating the immune response and therapeutic efficacy in cancer^19^, and IDO1 mediates immune suppression in the tumor microenvironment^20^. This study assesses the relationship between IDO1 and B cells in preclinical cancer models.

## Materials and Methods

### Mice and cell lines

C57BL/6 mice (The Jackson Laboratory) were housed in a pathogen-free environment, and experimental protocols were approved by the Johns Hopkins Institutional Animal Care and Use Committee. Similar to others studying the role of IDO1 in cancer^21,22^, female mice were used. We used young adult (ages 2-6 months) mice that were matched for age within each experiment. Experiments were performed at least twice with three to nine mice per group per experiment, and data from each experimental replicate were pooled for final analyses. The exception was Fig. 1G which was performed once with seven mice per group.

**Figure 1.**
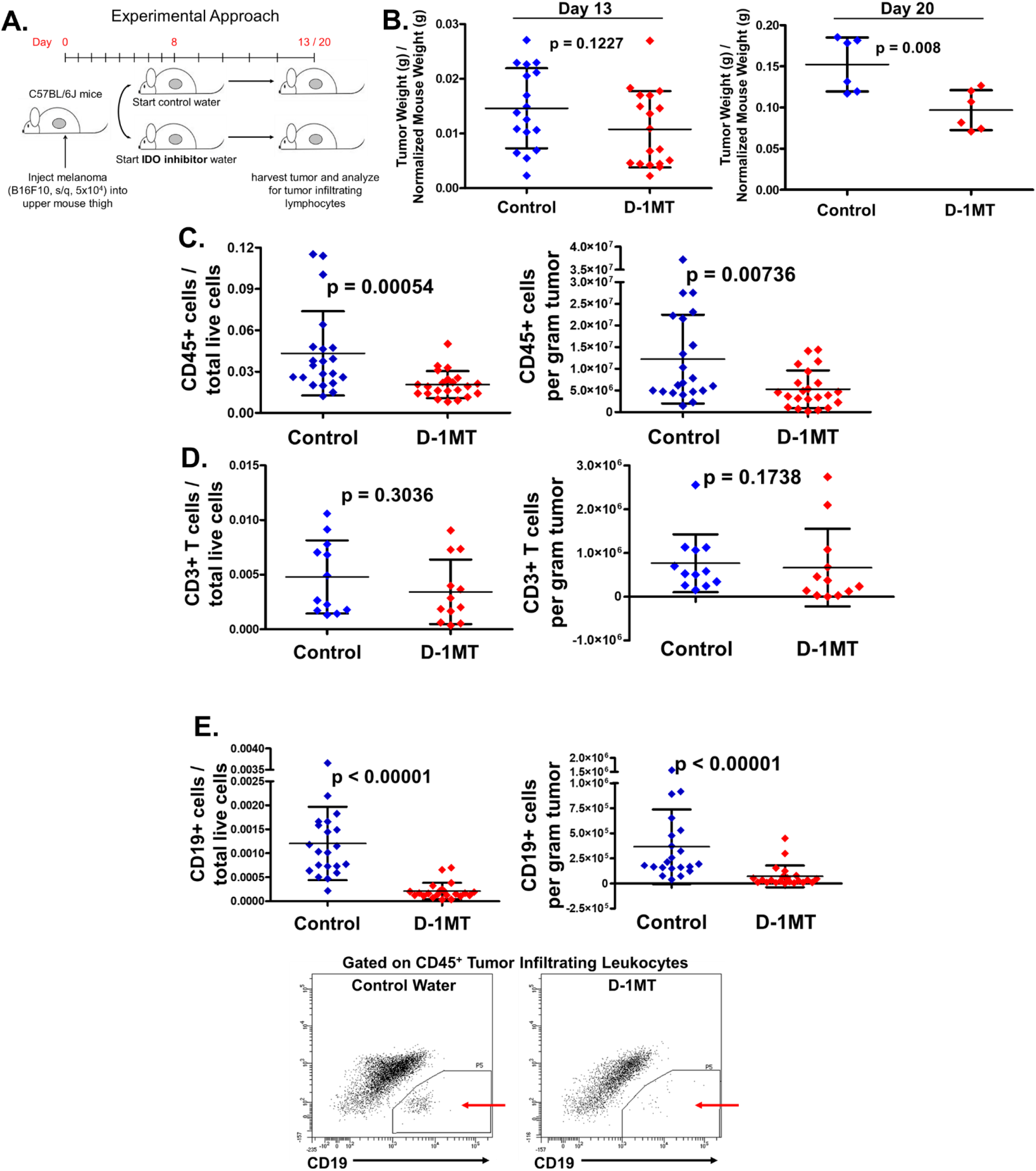
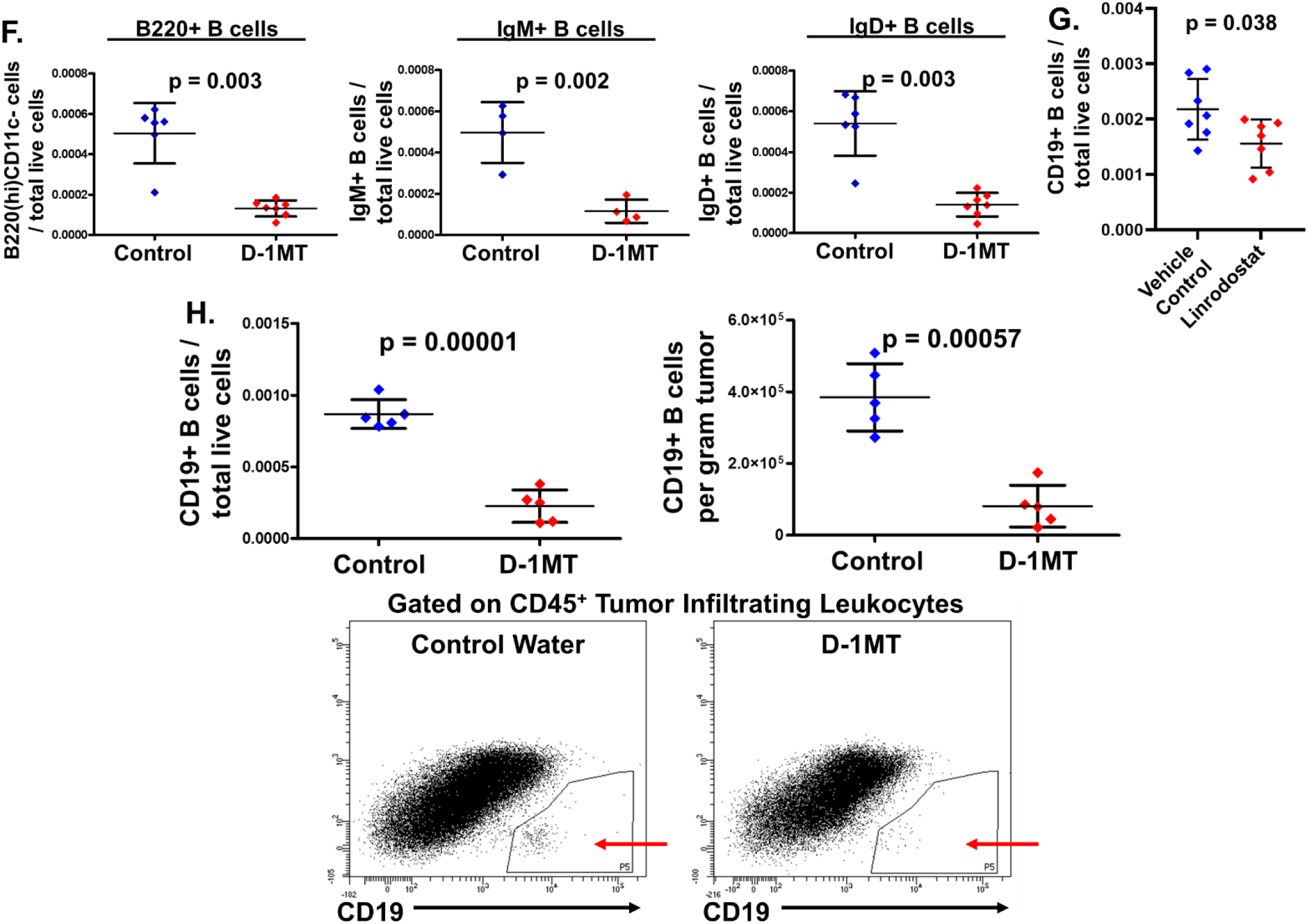
The IDO1 pathway regulates TIL-B cells. A. Treatment schema. D-1MT or control water was started on day 8, and tumors were harvested for flow analysis on day 13. B. B16F10 tumor weights at day 13 and 20. For day 20 harvest, mice received control or D-1MT water for 5 continuous days followed by 2 days of standard water, in order to minimize previously described dehydration that occurs with D-1MT^72^ and similar to other protocols that give a 2 day break after 5 days of D-1MT treatment^24^. C-E. B16F10 tumors were harvested on day 13 and analyzed for (C.) CD45^pos^ leukocytes, (D.) CD45^pos^CD3^pos^ T cells, or (E.) CD45^pos^CD19^pos^ TIL-B cells (with representative flow dot plots of TIL-B cells). F. Melanoma infiltrating CD45^pos^ cells from day 13 tumors were analyzed for B220^pos^CD11c^neg^ (left), IgM^pos^ (center), and IgD^pos^ (right) B cells. G. B16F10 tumors were treated as in (A.) except on day 8 vehicle control or linrodostat was started daily via oral gavage. Day 13 tumors were analyzed for CD45^pos^CD19^pos^ TIL-B cells. H. LLC tumors were injected and harvested as in (A.) and analyzed for CD45^pos^CD19^pos^ B cells. Each data point refers to a single mouse, and error bars represent ± standard deviation.

B16F10 mouse melanoma^23–25^ and Lewis Lung Carcinoma (LLC)^21,26^ were cultured in RPMI 1640 media with 10% Fetal Bovine Serum (FBS). All cell lines were propagated in flasks at 37°C and 5% CO_2_. By STR authentication on November 6, 2019, B16F10 and LLC cell lines were found to match the established genetic profile.

### Tumor Injections and D-1-methyl-tryptophan

For solid tumor models, we injected 5×10^4^ B16F10 cells or 1×10^5^ LLC cells (diluted in sterile PBS) subcutaneously into the lower left mouse thigh (considered day 0). For experiments with flow cytometry, the IDO1 pathway inhibitor D-1-methyl-tryptophan (D-1MT or indoximod, 2mg/mL, Sigma) or control saccharin water was given to mice in bottles protected from light beginning on day 8, with tumor, spleen, and/or bone marrow harvest on day 13. For experiments with the direct IDO1 inhibitor linrodostat (BMS-986205, Selleck Chemicals), mice were treated with vehicle (200 µL of 40 proof ethanol), or 100 mg/kg linrodostat (diluted in sterile water from a stock solution of 200 proof ethanol to 40 proof ethanol just before gavage) via oral gavage daily. Treatments were started on day 8, with harvest on day 13.

For experiments involving MDSC depletion, anti-Gr1 antibody (Biolegend, Cat. No. 108453) or isotype control antibody (Biolegend, Cat. No. 400671) was administered (200 µg, intraperitoneal) on day 7, 9, and 12 following injection with B16F10 melanoma on day 0. Mice were harvested on day 13.

For tumor growth experiments, control or D-1MT water was initiated on day 8. Tumors were measured using calipers every 3-4 days, and tumor volume was calculated using the formula V = L*(W^2^)*(π/6), where V is the tumor size in mm^3^, L is the longest diameter, and W is the shortest diameter^27^. Mice were weighed at baseline and with each tumor measurement.

### Preparation for Tumor Infiltrating Lymphocyte Analysis

After mouse sacrifice on day 13 (or day 20 in **Fig. 1B** as noted), tumors were isolated and weighed. Each tumor was placed in a separate well in a 6 well plate in RPMI media supplemented with 10% FBS, 1000 units/mL collagenase, and 0.05 mM β-mercaptoethanol. Tumors were dissected manually, followed by incubation for 1 hour at 37°C. Cells were filtered through a 70 µm cell strainer and resuspended to a total concentration of 2×10^7^/mL before staining. For spleen and bone marrow analysis, red cells were lysed with ACK (Ammonium-Chloride-Potassium) Lysis Buffer (ThermoFisher) before counting and staining.

### Analytical Flow Cytometry

We used the following antibodies for cell surface staining: CD19 (clone 6D5), Gr-1 (RB6-8C5), IgD (11-26c.2a), PD-1 (29F.1A12), PD-L1 (10F.9G2), PD-L2 (TY25), CD21/35 (7E9), CD23 (B3B4), CD24 (30-F1), Tim1 (RMT1-4), CD5 (53-7.3), and CD25 (PC61; all from Biolegend), and CD45 (clone 30-F11), CD19 (1D3), CD11b (M1/70), B220 (RA3-6B2), IgM (R6-60.2), and CD1d (1B1; all from BD Biosciences). Cells were first incubated with CD16/CD32 Fc block (BD Biosciences), followed by a cell viability stain (either Live/Dead Fixable Aqua Dead Cell Stain Kit, 405 nm excitation, Invitrogen ref. no. L34966; or Fixable Viability Stain 700; BD Horizon Cat. No. 564997). For tumor cell analysis, controls containing only viability stain were used to set initial gating structure, to exclude dead cells and doublets (**Sup. Fig. 1**). Fluorescence minus one controls were used as in **Sup. Fig. 1**, to determine cells staining positive for a particular fluorophore above background, as described^28,29^. Leukocytes were distinguished from tumor cells by CD45 staining. The LSR II flow cytometer or FACSCelesta (BD Biosciences) was used for cell acquisition, and Diva software was used for analysis.

### RNA Sequencing Analyses

All code for data acquisition, preprocessing, analysis, and figure generation is available on GitHub. TCGA bulk RNA sequencing data for muscle invasive bladder cancer, lung adenocarcinoma, squamous cell lung carcinoma, and melanoma were downloaded from the Genomic Data Commons using the GenomicDataCommons package^30^. Single cell RNA sequencing (scRNAseq) data for implanted B16F10 tumors are publicly available^31^ and were downloaded from ArrayExpress: E-MTAB-7427.

For pre-processing of scRNAseq data, TPM (transcripts-per-million) expression was downloaded, data were pre-filtered, and expression was scaled by log2(TPM + 1). TCGA data were normalized and scaled using DESeq2’s vst (variance stabilizing transformation)^32^.

For data analysis, GSVA^33^ (geneset variation analysis) was performed using a Gaussian kernel estimation of the cumulative distribution function. Scores were calculated using the mx.diff mode such that under the null hypothesis the distribution of scores would approximate a normal distribution with mean 0. Scores that were above 0 were labelled as ‘hi’, while those below zero were labelled as ‘lo’. For gene expression, a cell is ‘+’ for a gene if the expression cutoff is > 0 but ‘hi’ if the expression of a gene is obviously bimodal, setting the cutoff between low/no expression and high expression. R Version 4.0.3 was used for all scRNAseq and bulk RNA sequencing analyses^34^.

### IDO-specific siRNA expression vectors

*Ido1* shRNAs targeting different specific sequences of *Ido1* (shRNA-Ido1-908 and shRNA-Ido1-909) expressed by a lentiviral vector were designed by the Broad Institute (Cambridge, MA, US) and purchased from MilliporeSigma (Cat No. SHCLNG-NM_008324; clone ID: TRCN0000066908; clone ID: TRCN0000066909). The empty shRNA vector, pLKO.1 (TRC) (MilliporeSigma, Cat No. SHC001) was used as a control.

### Lentivirus Production

For lentivirus production^35,36^, COS1 African green monkey kidney epithelial cells (American Type Culture Collection, Cat. No. ATCC^®^ CRL1650™) were transfected with pCMV-ΔR8.91 (Creative Biogene, Cat. No. OVT2971) and pCMV-VSVg packaging plasmids (Addgene, Cat. No. 8454) and the *Ido1* shRNA using Lipofectamine 2000 (ThermoFisher Scientific, Cat. No. 11668027). Cells were transfected in virus collection medium (DMEM supplemented with 1% FBS) which was replaced with fresh medium 6 h after transfection. The culture supernatants were collected at 24h, 48h and 72h after transfection, filtered, and concentrated.

### Generation of B16F10 IDO1 silenced stable cell lines

B16F10 cells were transduced using standard lentivirus transduction protocols (see Addgene protocol found at: http://www.addgene.org/tools/protocols/plko/#E). Briefly, B16F10 cells were seeded into 24-well plates and grown to ∼80% confluence then each of the 3 viruses (TRC, shRNA-Ido1-908 and shRNA-Ido1-909) plus a mock control were added in triplicate wells in complete medium (DMEM, 10% FBS) supplemented with 100 U/ml penicillin, 100 ug/ml streptomycin (ThermoFisher Scientific, Cat. No. 15140-122) plus 8 ug/ml polybrene transfection reagent (MilliporeSigma, Cat. No. TR-1003-G). Cells were incubated at 4-8 °C for two hours then placed in a 37 °C incubator overnight. Cells were then split into 6-well plates in complete medium and incubated for two days to allow recovery before placing under 2 ug/ml puromycin selection (MilliporeSigma, Cat. No. P8833-25MG).

### Analysis of Ido1 silencing in transduced B16F10

B16F10 cells normally have very low levels of *Ido1* expression. Therefore, in order to test the potency and efficacy of the *Ido1* silencing, we induced *Ido1* by incubating 200 U/ml recombinant murine IFN-γ (PeproTech, Cat. No. 315-05) at 37°C for 5 hours prior to harvesting for RNA expression by qRT-PCR. Total RNA was extracted from the parental and *Ido1* silenced B16F10 cell lines using the Qiashredder (Qiagen, Cat. No. 79654) and RNeasy mini kit (Qiagen, Cat. No. 74104) with on-column RNase-Free DNase digestion (Qiagen, Cat. No.79254). First strand cDNA was generated using the High-Capacity cDNA Reverse Transcription Kit (ThermoFisher Cat. No. 4368814). Real-time PCR was performed on an AriaMx Real-Time PCR system (Agilent) using primer/probes from ThermoFisher targeting exon 5 of the *Ido1* gene (Mm00492590_m1) and normalized to beta-actin (P/N 4352933E). Amplification was performed using TaqMan Universal PCR Master Mix (ThermoFisher Scientific, Cat. No. 4324018) and analyzed using the ^ΔΔ^Ct method^37^.

### Statistics

For mouse experiments, data from each experimental group was tested for normality by the Ryan-Joiner Test^38^ and D’Agostino-Pearson Test^39^. The student’s t-test was used if data were normally distributed; the Mann-Whitney test was used otherwise^40^. P values ≤ 0.05 were considered significant. For scRNAseq and bulk RNA sequencing analyses, testing for equivalence of means was performed using a two-tailed Welch’s unequal variances t-test, with a significance cutoff of p < 0.05. As only a single comparison was made for each dataset, no multiple testing was applied.

## Results

### IDO1 pathway inhibition decreases tumor infiltrating B cells

To explore the role of the IDO1 pathway in antitumor immunity, we used D-1-methyl-tryptophan (D-1MT, indoximod), a tryptophan mimetic with established pharmacodynamics in humans and mice^41,42^, in the well-established implantable B16F10 model, which is an aggressive melanoma cell line that is wild-type for *BRAF*. Eight days after implanting B16F10 melanoma tumor cells into flanks of immunocompetent mice (see **Fig. 1A**)^23,24^, mice were treated with either D-1MT administered in drinking water or control water as previously described^47^. Similar to previous reports, IDO1 pathway inhibition had no significant effect on tumor size within the first 5 days of therapy (day 13 after implantation), although there was a modest decrease in tumor size after 12 days of therapy (day 20 after tumor cell implantation; **Fig. 1B**). To define the effects of D-1MT on the immune landscape, we first examined tumor infiltrating leukocytes (TILs) at early stages after tumor implantation (day 5 of therapy) using flow cytometry. There was a modest, but consistent, decrease in TIL frequency and absolute number per gram of tumor tissue (**Fig. 1C**). To determine whether intra-tumoral T cells changed with D-1MT treatment, we used the pan T cell marker CD3. Similar to prior results in a syngeneic murine model of lung cancer^22^, we found no change in CD3^pos^ T cell frequency or absolute number with D-1MT treatment (**Fig. 1D**).

Because IDO1 regulates B cells in non-cancer models^18,48^, we tested whether D-1MT affected the frequency and/or absolute number of TIL-B cells using the pan B cell marker, CD19. In control, untreated mice, the frequency and number of TIL-B cells is rare, comprising only ∼0.1% of total live cells (**Fig. 1E**). Strikingly, however, there was a profound decrease in CD19^pos^ TIL-B cell frequency and absolute number per gram tumor in D-1MT treated mice after 5 days of treatment (**Fig. 1E**), suggesting that D-1MT treatment either decreased the total B cell number and/or expression of CD19 on the B cell surface. To help distinguish between these possibilities, we tested whether TILs expressing other pan-B cell markers were also decreased, reasoning that a reduction would favor a decrease in total B cell number, since it would be less likely for D-1MT to repress expression of multiple B cell markers. To this end, we included B220 (a marker for B cell precursors and mature B cells), IgM (a marker for both immature and mature B cells), and IgD (found on late transitional and mature B cells). TIL-B cells expressing these markers decreased following IDO1-inhibitor therapy, consistent with a decrease in B cell frequency and number (**Fig. 1F**).

To determine whether D-1MT mediated decrease in TIL-B cells was due to an on-target effect of IDO1 inhibition, we treated melanoma bearing mice with the direct IDO1-inhibitor linrodostat. Consistent with the above results, linrodostat treatment significantly decreased TIL-B cells (**Fig. 1G**). To determine whether these results are generalizable to another tumor model, we tested a murine lung cancer implantation model (LLC)^21,26^. Similar to our findings in melanoma, D-1MT treatment significantly decreased TIL-B cell infiltration in the LLC model (**Fig. 1H**). Together, these data suggest that IDO1 pathway inhibition decreases the number of TIL-B cells in preclinical models of melanoma and lung cancer.

### Melanoma TIL-B cells have a heterogeneous phenotype

Recent data suggests improved prognosis in melanoma patients with high levels of TIL-B cells^49^, however increases in TIL-B cells also seems to correlate with resistance to *BRAF* targeted therapy^50^. To more fully understand this dichotomy, we sought to characterize this population further by analyzing TIL-B cells from publicly available single cell RNA sequencing (scRNAseq) data from implanted B16F10 tumors^31^. We found that TIL-B cells were uniformly positive for *Ighm*, and most expressed *Ighd*, suggesting TIL-B cells are naïve or transitional (**Fig. 2A, left**). Next, we determined that TIL-B cells did not experience class switching, as TIL-B cells did not express IgG or IgA heavy chain genes, further indicating that TIL-B cells are naïve or transitional. TIL-B cells also express IgM, IgD, and the pan-B cell marker B220 by flow cytometry (**Sup. Fig. 2A**). We found that 19% of TIL-B cells express the transitional B cell specific gene *Cd93*^51^ (**Fig. 2B, left**), 44% of TIL-B cells express the marginal zone specific gene *Mzb1* (**Fig. 2B, center**) and 62% of TIL-B cells express *Cd27* (**Fig. 2B, right**), an activation marker found on B1 or innate memory cells (*CD27*^*pos*^*IghD*^*pos*^), or naïve or transitional cells (*CD27*^*neg*^*IghD*^*pos*^) (**Sup. Fig. 2B**).

**Figure 2.**
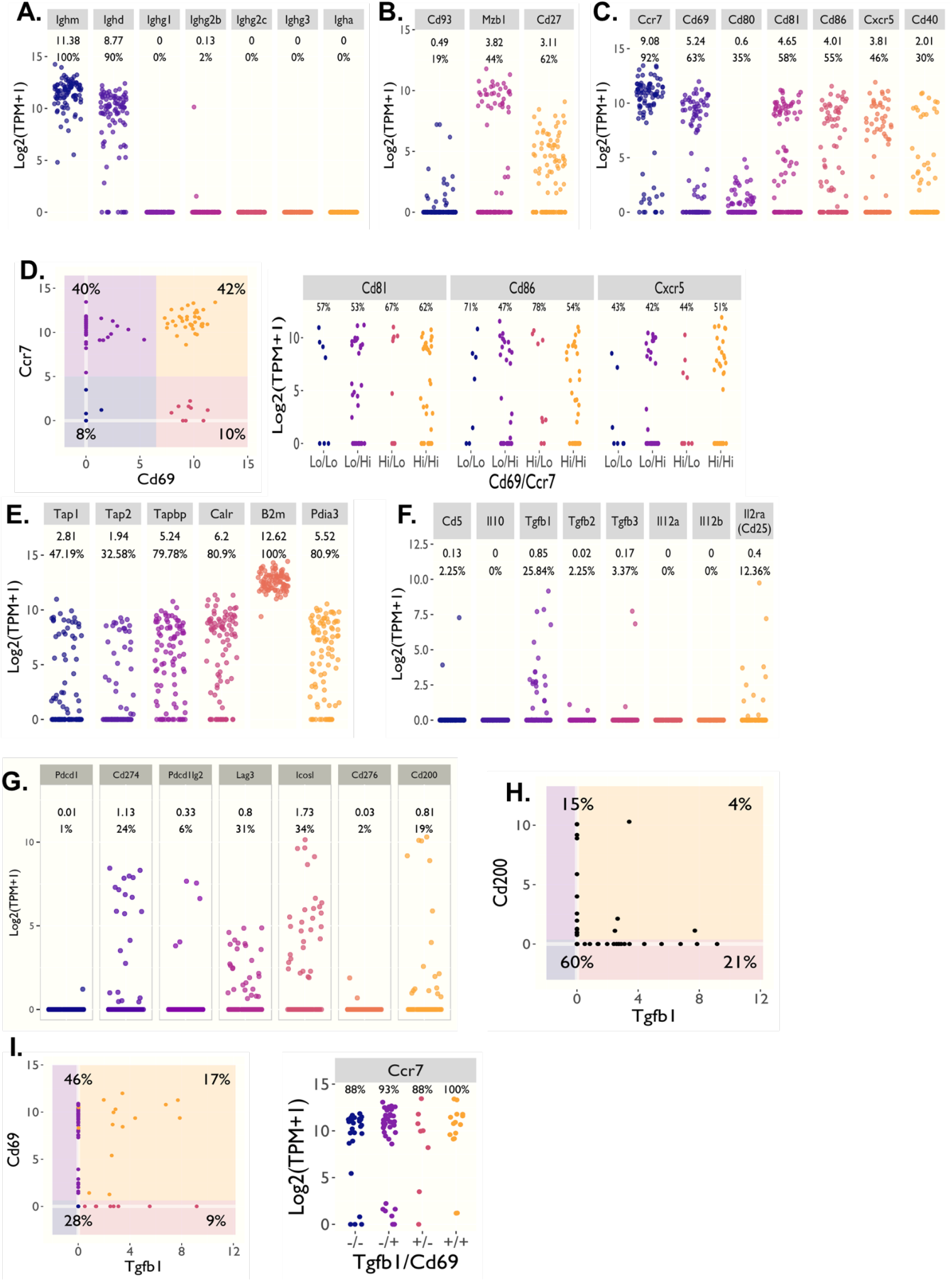
Melanoma TIL-B cells are phenotypically and functionally heterogeneous. ScRNAseq data from B16F10 tumors was accessed from a publicly available dataset^31^ and TIL-B cells analyzed for the following phenotypic and functional genes: A. Surface immunoglobulin genes, B. *Cd93* (left), *Mzb1* (center), and *Cd27* (right). C. Genes associated with B cell activation. D. TIL-B cells were graphed based on expression of *Ccr7* and *Cd69* (left), and clusters based on *Cd69 / Ccr7* expression were graphed based on *Cd81* (left), *Cd86* (center), and *Cxcr5* expression (right). E-G. TIL-B cells were graphed based on expression of (E.) antigen processing genes, (F.) immune suppressive genes, and (G.) genes encoding checkpoint molecules. H. TIL-B cells were plotted based on expression of *Tgfb1* and *Cd200*. I. TIL-B cells were plotted based on *Tgfb1* and *Cd69* expression, and clusters based on *Tgfb1 / Cd69* expression colored based on *Ccr7* expression.

To define the function of TIL-B cells, we first examined TIL-B cells for expression of genes encoding activation markers and costimulatory molecules. Most TIL-B cells express *Ccr7* (**Fig. 2C**), which is expressed by a TIL-B cell subset that correlates with response to checkpoint immunotherapy in patients with advanced melanoma^19^. A majority of TIL-B cells express genes encoding the activation markers CD69, CD81, and CD86, but fewer TIL-B cells express genes encoding for CD80 (35%), the costimulatory molecule CD40 (30%), and the chemokine receptor CXCR5 (46%), which facilitates B cell receptor mediated activation^52^ (**Fig. 2C**).

To examine co-expression of activation markers by TIL-B cells, we graphed *Ccr7* and *Cd69* along with other activation related genes. Most TIL-B cells are *Ccr7*^*high*^ *Cd69*^*high*^ (42% of all TIL-B cells) or *Ccr7*^*high*^ *Cd69*^*low/neg*^ (another 40%) (**Fig. 2D, left**). About half of *Ccr7*^*high*^ *Cd69*^*high*^ and *Ccr7*^*high*^ *Cd69*^*low/neg*^ TIL-B cells express *Cd81, Cd86*, and/or *Cxcr5* (**Fig. 2D, right**), suggesting many TIL-B cells are in an activated state. Most TIL-B cells express antigen processing (**Fig. 2E**) and major histocompatibility complex (MHC) genes (**Sup. Fig. 2C**), suggesting many TIL-B cells are primed for antigen presentation.

Next, we interrogated the scRNAseq data for expression of genes associated with immune suppression. TIL-B cells do not express genes found in a subset of Bregs called B10 cells^53^ (*Cd5* or *Il10*, **Fig. 2F**). Moreover, TIL-B cells do not express the IL-35 associated gene *Il12a*, expressed by IL-35 expressing Bregs^54^ (**Fig. 2F**). However, consistent with previous reports that TGFβ is produced by a Breg subset^55^, ∼26% of TIL-B cells express *Tgfb1* (**Fig. 2F**). Additionally, a subset of TIL-B cells express *Il2ra* (CD25), which identifies a Breg subset^56^ (**Fig. 2F**).

Finally, we interrogated TIL-B cells for expression of immune checkpoints targeted by commonly used immunotherapeutic agents. Consistent with recent data demonstrating TIL-B cells promote anti-melanoma immunity in response to PD-directed therapy^19^, TIL-B cells express *Cd274* (PD-L1), but only minimal *Pdcd1* (PD-1) or *Pdcd1lg2* (PD-L2) (**Fig. 2G**). A minority of TIL-B cells express *Lag3*, and very few TIL-B cells express the inhibitory costimulatory molecule B7-H3 (*Cd276*, **Fig. 2G**). However, a significant minority of TIL-B cells express the inhibitory checkpoint *Cd200* (**Fig. 2G)**.

To determine whether TIL-B cells co-express immune inhibitory genes, we graphed TIL-B co-expression of *Tgfb1* and *Cd200* (**Fig. 2H**). Very few TIL-B cells co-expressed *Tgfb1* and *Cd200* (**Fig. 2H**). As one report suggests Bregs can express *Cd69*, we graphed *Cd69* versus *Tgfb1* expression (along with *Ccr7*) and found a fraction of TIL-B cells (∼15%) were *Tgfb1*^*pos*^*Cd69*^*high*^*Ccr7*^*high*^ (**Fig. 2I**). Most TIL-B cells expressing multiple activation markers are likely immune stimulatory, however a cohort of TIL-B cells within this subset may have inhibitory function.

In summary, most CD19^pos^ TIL-B cells express *Ighm* and *Ighd*, and are not class-switched. TIL-B cells are markedly heterogeneous, including transitional and marginal zone phenotypes. While a minority of TIL-B cells express immune suppressive genes, most have an activated phenotype.

### IDO1 pathway inhibition decreases splenic transitional B cells

Clinical studies have shown that patients treated with IDO1-inhibitor drugs can develop a transient lymphopenia^42^, and our studies demonstrate D-1MT decreases TIL-B cells in mice. To determine whether the effect of D-1MT on TIL-B cells results from a systemic decrease in B cells, we analyzed spleens from control and D-1MT treated tumor-bearing mice. There was a significant decrease in total splenic B cells in mice with B16F10 tumors (**Fig. 3A**) and LLC tumors (**Fig. 3B**) after D-1MT treatment. D-1MT treatment also decreased spleen weights (normalized to mouse weight) in tumor-bearing mice, consistent with the decrease in total intrasplenic B cells (**Sup. Fig. 3**). Because IDO1 is known to be inducible in a counter-regulatory fashion in settings of inflammation (including tumors)^45,57^, we hypothesized that D-1MT treatment in mice lacking tumors would not significantly reduce splenic B cells, which is the result we observed (**Fig. 3C**).

**Figure 3.**
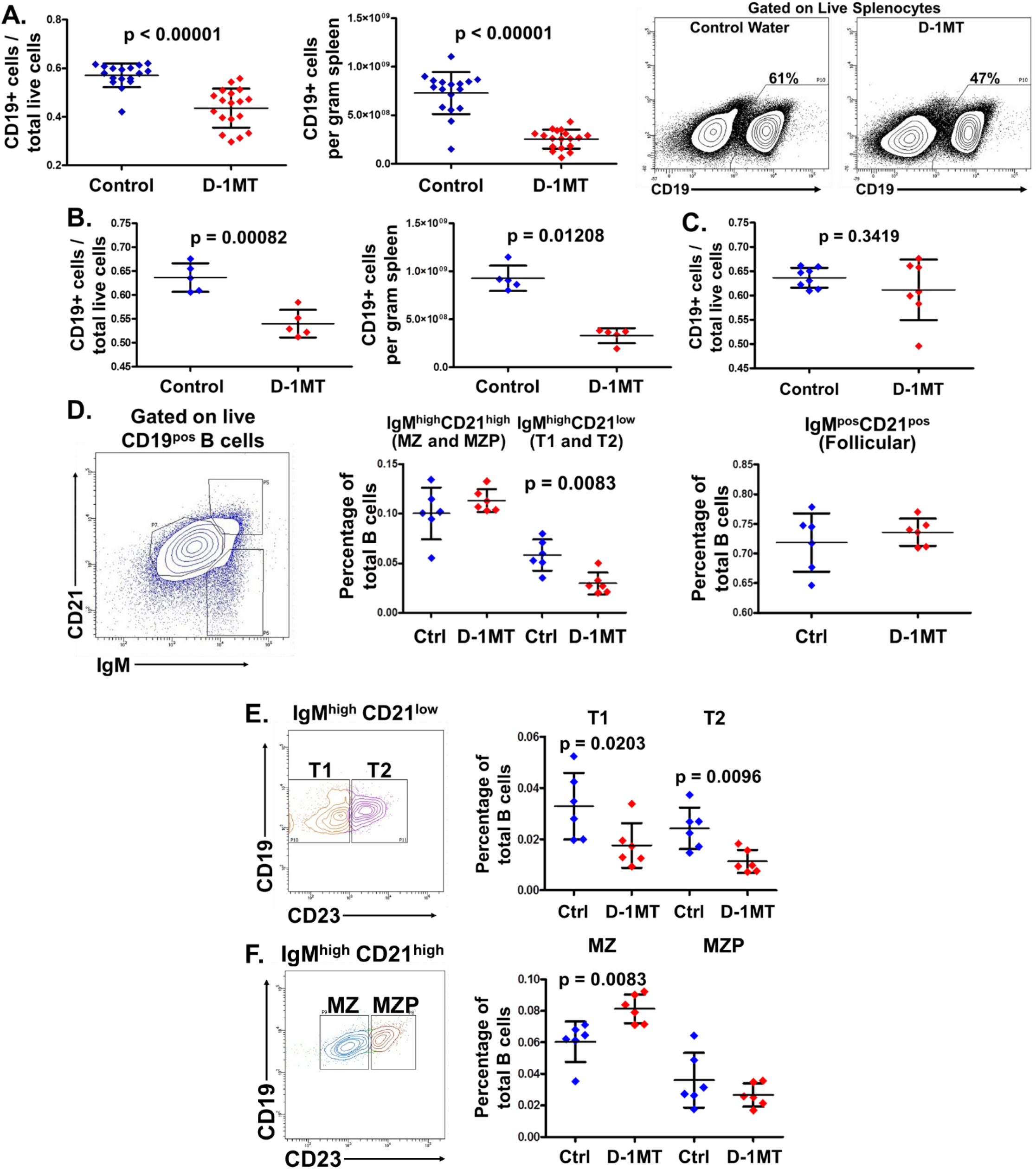
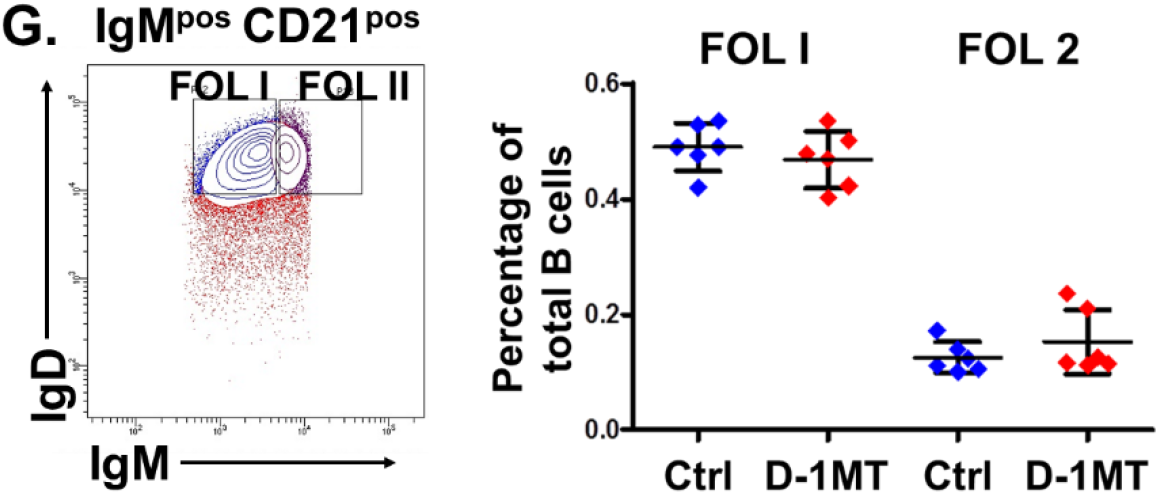
The IDO1 pathway regulates transitional B cells in melanoma bearing mice. Melanoma (B16F10) or Lewis Lung Carcinoma (LLC) cells were implanted into wild-type mice as in **Fig. 1** (except Fig. 3C) and treated with control water or D-1MT until harvest on day 13. A-B. CD45^pos^CD19^pos^ B cells from day 13 spleen of (A.) B16F10-bearing, (B.) LLC-bearing, or (C.) non-tumor-bearing mice were analyzed by flow cytometry. D. Splenic B cells from B16F10-bearing mice were divided into IgM^high^CD21^high^ (containing marginal zone B cells, MZ and marginal zone precursors, MZP), IgM^high^CD21^low^ (containing transitional B cells), and IgM^pos^CD21^pos^ (containing follicular B cells). E-G. Splenic B cells from the subdivisions described in (D.) were further divided into transitional (T)1 and T2 B cells (E.), MZ and MZP B cells (F.), and follicular (FOL) I and FOL II B cells (G.).

In order to determine whether D-1MT impacts the same B cell subset within tumors and spleen, we performed additional phenotypic analysis of splenic B cells. To do this, we initially classified splenic CD19^pos^ B cells into marginal zone plus marginal zone precursors (IgM^high^CD21^high^), transitional (IgM^high^CD21^low^), and follicular (IgM^pos^CD21^pos^) subsets. We found that D-1MT decreased splenic transitional B cells, but not follicular B cells (**Fig. 3D**). In order to further subdivide B cells, we used CD23 co-staining to subdivide marginal zone and follicular B cells, and IgD to subdivide follicular B cells. D-1MT decreased both transitional 1 (T1) and transitional 2 (T2) B cell subsets (**Fig. 3E**), with an increase in marginal zone B cells (**Fig. 3F**). Conversely, D-1MT did not impact splenic follicular or marginal zone precursor B cell numbers (**Fig. 3F-G**). Thus, D-1MT decreased the splenic transitional B cell pool, which suggested that D-1MT impacted TIL-B cells via a systemic decrease of B cells.

### IDO1 pathway inhibition leads to a partial block in B cell development

To determine how D-1MT decreased B cells systemically, we investigated its effects on B cell development within the bone marrow in tumor bearing mice. There was a marked decrease in B cell precursors (B220^pos^IgM^neg^) in the bone marrow (**Fig. 4A-B**). There was also a decrease in pro-B and pre-B cells, together with an enrichment in the immature prepro-B cell compartment within the total B cell precursor pool (**Fig. 4D-E**, schematic of B cell development in **Fig. 4C**). Consistent with this, D-1MT increased the ratio of prepro-B cells relative to more mature B cell precursors (**Fig. 4F**). Together, these data demonstrate that treatment with D-1MT decreases pro-B and pre-B cells within the bone marrow, suggesting that D-1MT imposes a partial block in B cell development.

**Figure 4.**
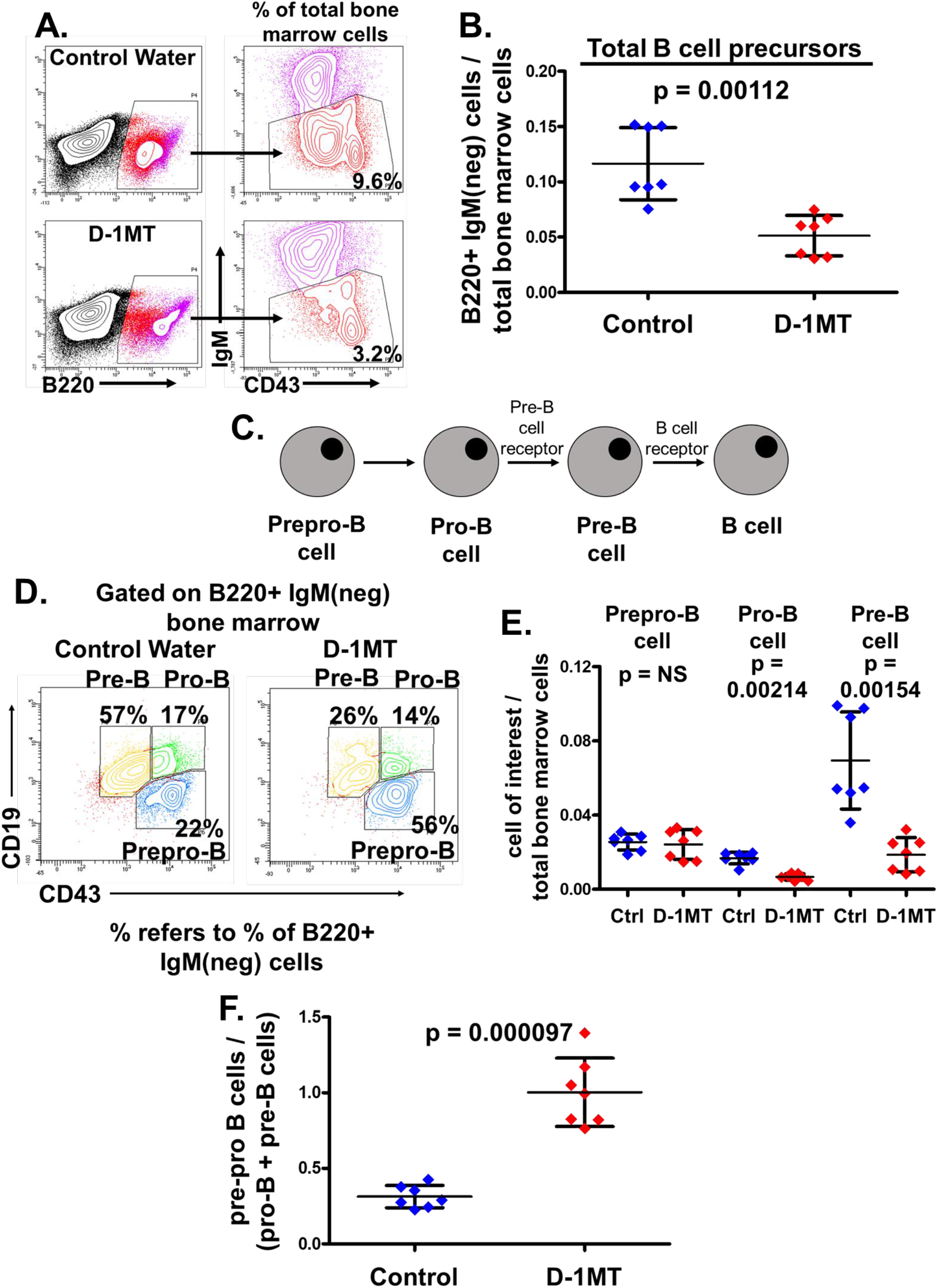
The IDO1 pathway regulates B cell precursors in melanoma-bearing mice. B16F10 tumors were implanted into wild-type mice as in **Fig. 1A** and treated with control water or D-1MT until harvest on day 13. A-B. Bone marrow from melanoma-bearing mice was harvested at day 13 and analyzed for B220^pos^IgM^neg^ B cell precursors. C. Schematic of B cell development. D. B220^pos^IgM^neg^ B cell precursors were analyzed for prepro-B, pro-B, and pre-B subsets as shown. E. Total bone marrow cells were analyzed for prepro-B, pro-B, and pre-B cells. F. Ratio of pre-pro B cells to downstream pro-B and pre-B cells. Bars represent standard deviation.

### D-1MT decreases intratumoral MDSCs, which regulate TIL-B cells

To dissect the role of IDO1 on other immune cells in the tumor microenvironment that may modulate TIL-B cells, we investigated MDSCs. MDSCs modulate B cell responses^58–60^; conversely, B cells modify MDSC function^61^, and MDSCs express *IDO1* in some settings^62–64^. Whether MDSCs are essential for TIL-B cell maintenance is unknown. Similar to B cells, we found that D-1MT decreases MDSCs (CD11b^pos^Gr-1^pos^) in mice bearing B16F10 (**Fig. 5A**) and LLC tumors (**Fig. 5B**). Further, D-1MT decreases splenic MDSCs in B16F10 (**Fig. 5C**) and LLC tumors (**Fig. 5D**). Because prior studies showed that MDSCs influence B cell function^59,65,66^, and D-1MT decreased both TIL-B cells and MDSCs, we sought to determine whether MDSCs are required for TIL-B cell maintenance in our model. To this end, MDSCs were depleted *in vivo*, using an anti-Gr1 antibody (**Fig. 5E**). Consistent with the known function of anti-Gr1 antibody, this treatment led to a significant decrease in both intratumoral and splenic MDSCs (**Sup. Fig. 4A-B**). Strikingly, systemic MDSC depletion resulted in a decrease in TIL-B cells (**Fig. 5F**), suggesting that MDSCs are required for TIL-B cell maintenance. Conversely, anti-Gr1 antibody increased the percentage of splenic B cells (**Fig. 5G**), unlike IDO1 pathway inhibition which decreased splenic B cells in melanoma bearing mice (**Fig. 3A**). Thus, the IDO1 pathway maintains intratumoral and systemic MDSCs, and MDSCs drive intratumoral (but not systemic) B cell maintenance.

**Figure 5.**
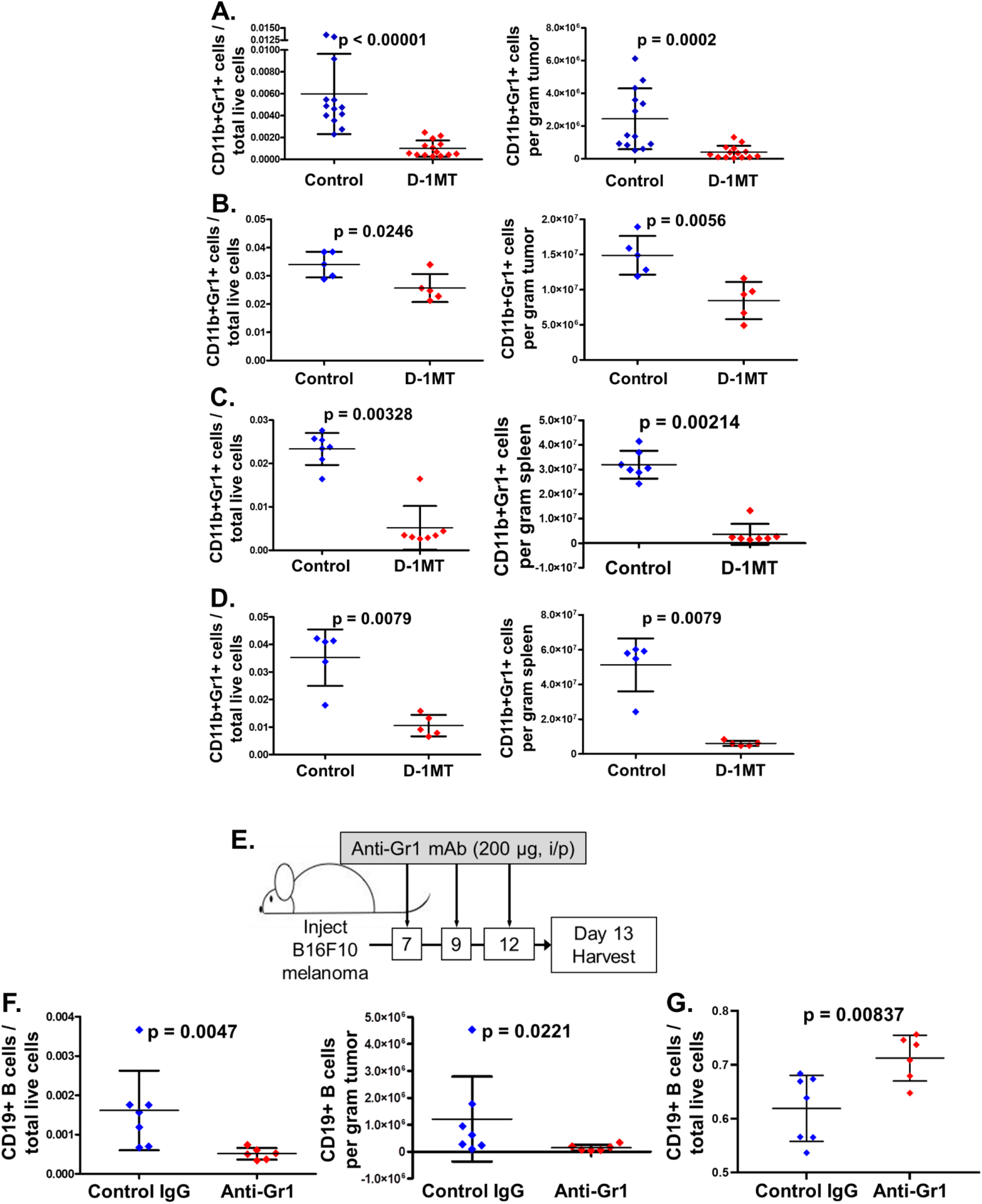
The IDO1 pathway regulates MDSCs, which regulate TIL-B cell maintenance. Mice were injected with B16F10 or LLC as in **Fig. 1**, and on day 13 tumors from B16F10 (A.) and LLC (B.) were analyzed for CD45^pos^CD11b^pos^Gr-1^pos^ MDSCs. C-D. Spleens from B16F10 (C.) and LLC (D.) tumor-bearing mice were analyzed for MDSCs. E. Treatment schema for MDSC depletion. Control Ig or Anti-Gr1 antibody (200 µg) was given via intraperitoneal injection on days 7, 9, and 12, with harvest on day 13. F. CD45^pos^CD19^pos^ TIL-B cells after MDSC depletion. G. Splenic CD19^pos^ B cells after MDSC depletion. Bars represent standard deviation.

### D-1MT treatment and tumor *Ido1* deletion have disparate immunologic impact

As D-1MT mediated modest effects on tumor growth with concomitant decrease in TIL-B cells and MDSCs, we studied whether these effects were due to tumor IDO1 via implantation of *Ido1* knockdown tumors. As *Ido1* is an inducible enzyme, we confirmed *Ido1* knockdown using interferon gamma stimulation *in vitro* (**Sup. Fig. 5)**. Similar to B16F10 tumors in mice treated with D-1MT, we found a modest (but significant) decrease in tumor size in *Ido1* knockdown tumors (**Fig. 6A**). We confirmed *Ido1* knockdown *in vivo* at time of tumor harvest (**Fig. 6B**). Unlike D-1MT treated B16F10 tumors, we found tumor specific *Ido1* deficiency led to a significant increase in CD45+ tumor infiltrating leukocytes (**Fig. 6C**) and no difference in CD19+ TIL-B cells (**Fig. 6D**), MDSCs (**Fig. 6E**), or T cells (**Fig. 6F**). Thus, while D-1MT and knockdown of tumor *Ido1* expression similarly impact tumor size, these interventions have disparate effects on B cells, T cells, and MDSCs.

**Figure 6.**
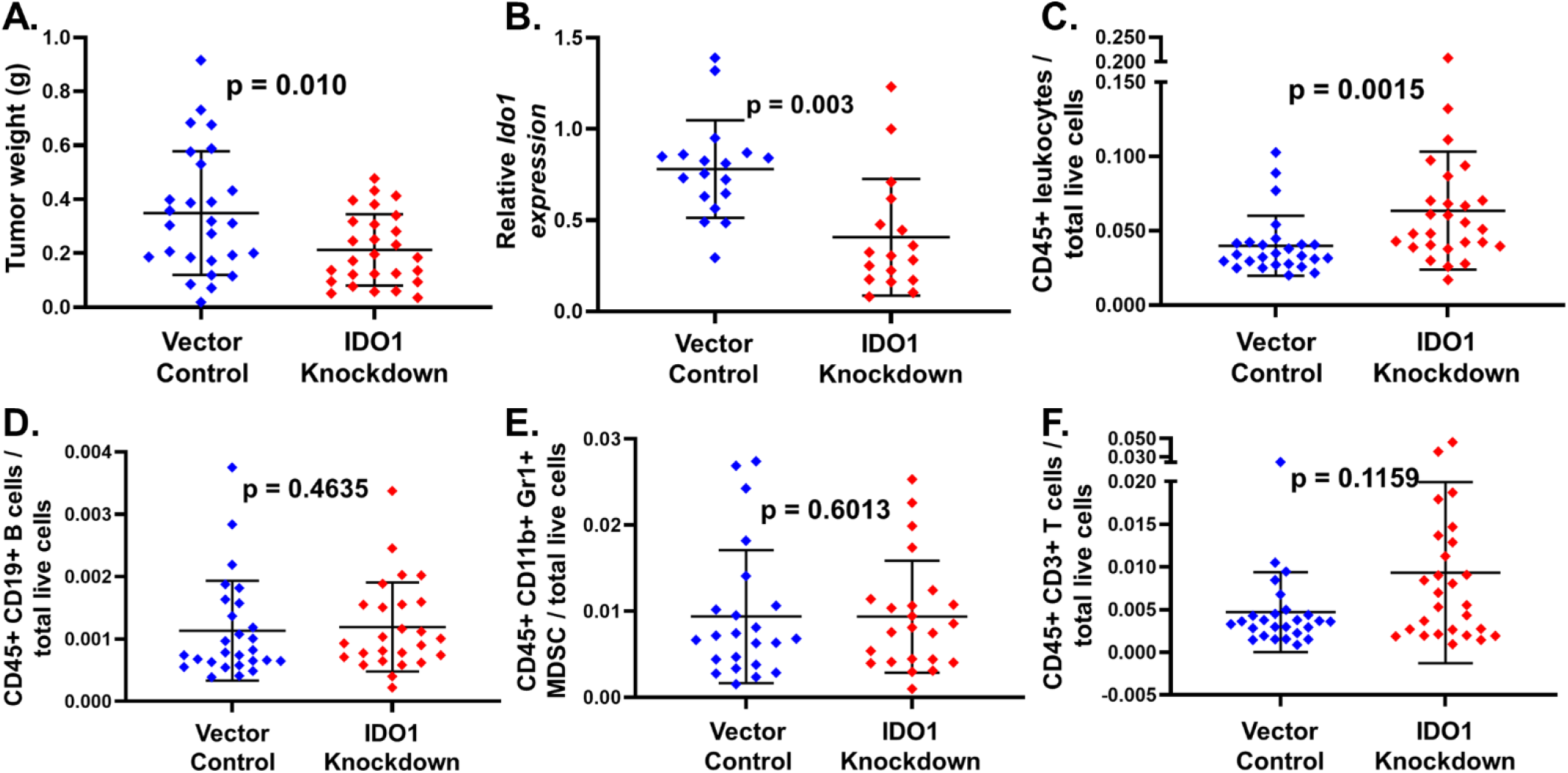
*Ido1* tumor knockdown impacts the tumor microenvironment differently than IDO1 pathway inhibition. Vector control or *Ido1* knockdown tumors were injected on day 0 and harvested on day 13. A. Tumor weights of vector control and *Ido1* knockdown tumors. B. *Ido1* expression of vector control and *Ido1* knockdown tumors *in vivo* following tumor harvest on day 13. C-F. Day 13 tumors were analyzed for (C.) CD45+ leukocytes, (D.) CD45+CD19+ B cells, (E.) CD45+CD11b+Gr1+ MDSCs, and (F.) CD45+CD11b+CD3+ T cells.

### *IDO1* is expressed by TIL-B cells

To determine whether IDO1 expression is associated with B cells in human cancers, we interrogated human tumors from The Cancer Genome Atlas (TCGA) for *IDO1* and B cell gene signature (BCGS) expression, using a validated BCGS^67^. We examined bladder cancer (BLCA), lung adenocarcinoma (LUAD), squamous cell carcinoma of the lung (LUSC), and melanoma (SKCM) and found each of these tumors segregated into two groups based on BCGS expression (low or high, **Fig. 7A**). Thus, we used the Partitioning Around Medoids (PAM) clustering algorithm to divide each tumor subtype into two groups based on high or low BCGS expression, and found intratumoral *IDO1* expression was significantly greater in tumors with high BCGS expression (**Fig. 7B**). To determine whether IDO1 is expressed by human TIL-B cells, we interrogated a publicly available dataset where tumors from melanoma patients were analyzed by scRNAseq^68^. Of all TIL-B cells in analyzed patient tumors, 377 of 515 (∼73%) expressed *IDO1* (**Fig. 7C**, left and **Fig. 7E**). However, using another dataset^69^, we found minimal (13 of 1379, or 0.94%) TIL-B cells expressed *IDO1* (**Fig. 7C**, right and **Fig. 7E**). To determine if this discrepancy was due to disparate sensitivity for *IDO1* detection, we plotted *IDO1* expression in all cells from analyzed tumors. While many cells (3246 of 4645, or ∼70%) express at least low levels of *IDO1* in the Tirosh dataset (**Fig. 7D**, left and **Fig. 7F**), comparatively few cells (278 of 16291, or 1.7%) express *IDO1* in the Sade-Feldman dataset (**Fig. 7D**, right and **Fig. 7F**), which primarily report cells that express high *IDO1*. The limited number of *IDO1* positive cells in TIL-B cells from the Sade-Feldman dataset may be due to technical limitations with detection of low expression of *IDO1*, which are robustly identified in the Tirosh dataset. Thus, TIL-B cells appear to express low levels of *IDO1* in patients with melanoma.

**Figure 7.**
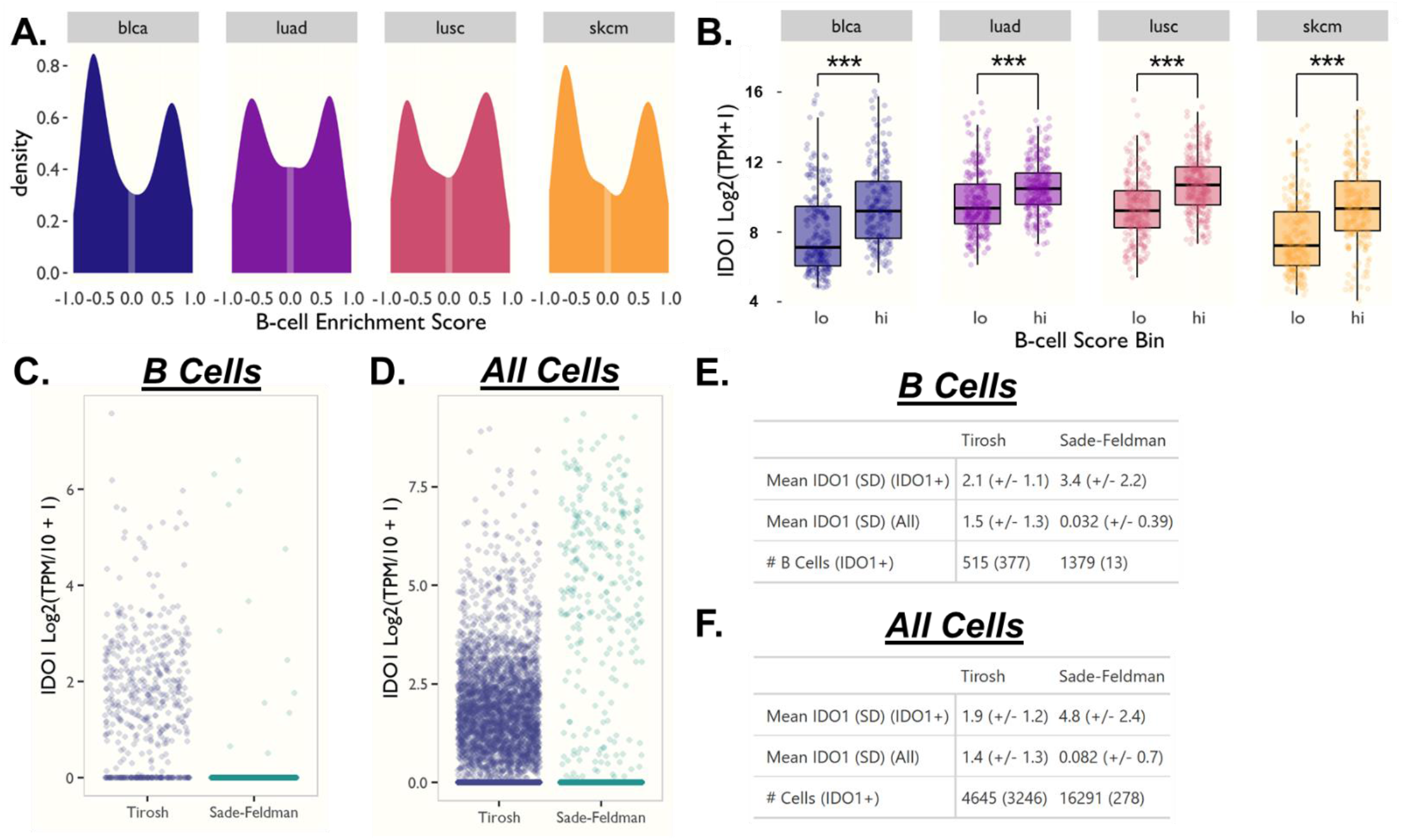
B cell gene signature (BCGS) expression correlates with *IDO1* in human cancers, and human intratumoral B cells express *IDO1*. A. Bulk RNA sequencing data from tumors from TCGA muscle invasive bladder cancer (blca), lung adenocarcinoma (luad), lung squamous cell carcinoma (lusc), and melanoma (skcm) were analyzed and graphed based on expression of a validated BCGS (Danaher et al.^67^). B. Tumors from (A.) were examined for correlation of *IDO1* with low and high levels of BCGS expression. *** = p < 0.001. C-D. Single cell RNA sequencing of melanoma tumors from Tirosh (left) and Sade-Feldman (right) datasets were analyzed for IDO1 expression (C. B cells, D. all cells). E-F. Results from C-D. are presented in table format.

## Discussion

IDO1 is considered a critical regulator of effector T cell responses in humans and mice^70–72^. IDO1 regulates innate and adaptive immune responses in settings that include maternal tolerance to paternal allo-antigens in pregnancy, control of autoimmunity, mucosal tolerance, chronic infection states, and others^70,73–75^. In cancer, the role of IDO1 in suppressing T cell anti-tumor immunity was strengthened by preclinical murine models^24,25,76^ and early phase clinical trials demonstrating efficacy of IDO1 inhibitors in combination with other agents^77,78^. Our study demonstrates a novel role for IDO1 in regulating the B cell compartment in cancer, both systematically and within established tumors.

The modest impact of IDO1 inhibition or tumor *Ido1* knockdown on tumor weight was expected, as the vast majority of preclinical and clinical studies support that IDO1-ablating treatments require combination with other therapies (chemotherapy, radiation, other immunotherapies) for significant anti-tumor effects^24,25,76,79^. Indeed, the best tumor response achieved in the phase I clinical trials using IDO1-inhibitor monotherapy was stable disease^42,80^.

The striking result in the current report is that IDO1 plays an important role in the regulation of systemic and intra-tumoral B cell subsets, and this effect may be mediated, at least in part, by MDSCs. We show for the first time that both D-1MT and linrodostat decrease intratumoral B cells. Recent studies have demonstrated that increased expression of a BCGS correlates with improved response to immune checkpoint inhibitor (ICI) therapy in patients with advanced melanoma^19^. Moreover, B cells are essential for ICI treatment to slow tumor growth in a mouse model of triple negative breast cancer, and this therapy is approved for use in humans^81,82^. Our analysis of scRNAseq data in a preclinical melanoma model suggests that most TIL-B cells in untreated tumors are stimulatory. This further suggests that combining IDO1 inhibition with ICIs may not be advantageous after all^83^.

Conversely, TIL-B cells are also associated with acquired resistance to BRAF-targeted therapy in melanoma^50^. Thus IDO1-inhibitor mediated TIL-B cell depletion may be beneficial in this setting, even as adjunctive therapy. Our data are also consistent with a report demonstrating IDO1 pathway inhibition attenuates pathology in a mouse model of rheumatoid arthritis, in part, by decreasing autoreactive B cells^84^. As metabolism of the amino acid glutamine is critical for B cell survival in certain conditions^16^, the role of IDO1 in regulating B cell development may be related to kynurenine depletion, and merits further study in rheumatologic conditions.

Our results demonstrating a decrease in MDSCs following pharmacologic IDO1 pathway inhibition is consistent with previous reports^62,85^. However, there is a discordance between genetic and pharmacologic models with both MDSCs and B cells, as genetic deletion of tumor *Ido1* does not impact tumor B cell or MDSC infiltration. Study of the IDO1-KO mouse in multiple settings, including pregnancy and implanted tumors, strongly suggest that these mice have developed compensatory mechanisms for the lack of IDO1 expression^24,86^. Consequently, robust reduction of tumor-infiltrating B cells and MDSCs via IDO1 inhibition may require IDO1 to first be upregulated in the tumor prior to IDO1-blockade. This is consistent with data that B cells and MDSCs can express IDO1^17,63^, and tumor *Ido1* over-expression increases TIL-MDSCs^87^.

Given that IDO1 appears to have a role in regulating MDSC tumor infiltration, it is remarkable that MDSCs themselves play a role in maintaining TIL-B cells. MDSCs can convert B cells into Bregs^60^ and block B cell proliferation and differentiation^59,65,66^. While this suggests that the absence of MDSCs could increase B cell numbers, we showed that MDSC depletion led to a *decrease* in TIL-B cells. Notably, MDSC depletion increased splenic B cells, suggesting MDSCs are not required to maintain systemic B cell populations. This is likely distinct from IDO1-mediated regulation of MDSCs, as IDO1 pathway inhibition decreased both TIL-B and splenic B cells. How MDSCs regulate TIL-B cells is an area of active study and may result in additional therapeutic targets for clinical investigations.

## Abbreviations

IDO1: indoleamine 2,3 dioxygenase-1
Bregs: regulatory B cells
Tregs: regulatory T cells
LLC: Lewis Lung Carcinoma
PD-1/L: programmed cell death protein-1/ligand
APCs: antigen presenting cells
MDSCs: myeloid derived suppressor cells
IL: interleukin

## Acknowledgements

The authors gratefully acknowledge support from the Conquer Cancer Foundation of the American Society of Clinical Oncology Young Investigator Award (to B.A.J.), the Harry J. Lloyd Charitable Trust Career Development Award (to B.A.J.), the AACR-BMS Fellowship for Young Investigators in Translational Immuno-oncology (to B.A.J.), the Alpha Omega Alpha Postgraduate Fellowship Award (to B.A.J.), two Johns Hopkins Physician Scientist Training Program Microgrant Awards (to B.A.J.), and the Johns Hopkins Institutional Training Grant in Medical Oncology (to B.A.J., NIH 5T32 CA009071, B.H. Park, PI). Initial studies were also supported by a NIH NRSA Predoctoral F30 award and an American Medical Association Foundation Seed Grant (both to B.A.J.). Further support was also provided by R01 CA235681 (to N.M.H.), and the Greenberg Bladder Cancer Institute. Dr. Johnson would also like to thank his patients, some of whom also graciously contributed financially to experiments published in this work.

## Dedication

This paper is dedicated to Ross C. Donehower, MD, Professor, Director of Medical Oncology, and Director of the Medical Oncology / Hematology Fellowship Program, Johns Hopkins School of Medicine, with gratitude for his superior mentorship and impact on B.A.J.’s career development.

## Author Contributions

B.A.J. conceived the study, designed, supervised, and executed experiments, analyzed data, and wrote the paper. K.A., J.M., W.Z., L.X., L.L., L.C., O.R., and D.W. performed experiments. D.D. and S.D. provided reagents/analytical tools. B.A.J., K.A., D.W., W.Z., L.L., L.X., L.C., O.R., D.D., S.D., T.S.J., N.H., D.J.M., and L.M.S.R. analyzed data. T.S.J. designed and performed pilot experiments. N.H., D.J.M., and T.S.J. revised the paper. L.M.S.R. supervised experiments, analyzed data, and wrote the paper.

---

**Supplemental Figure 1.**
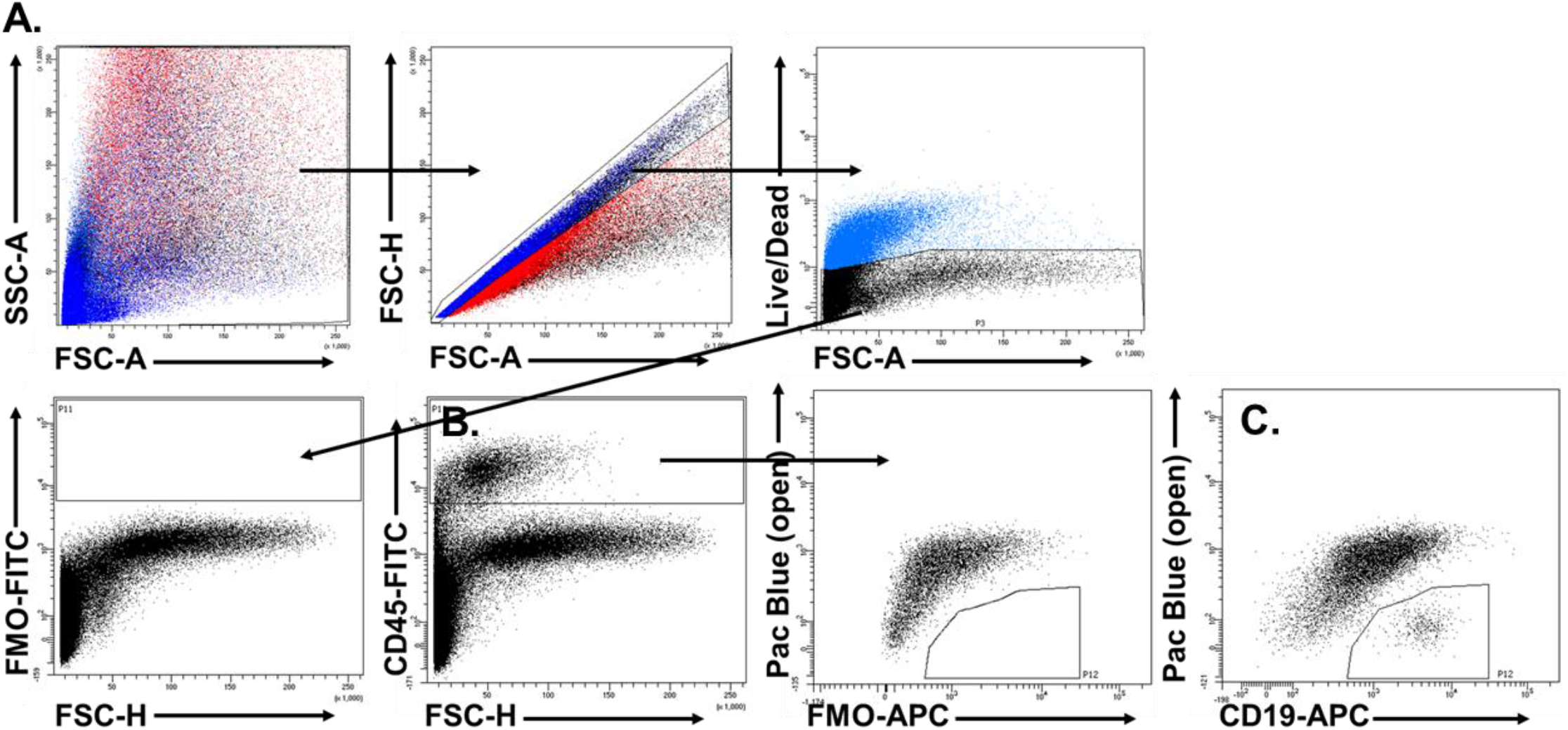
Gating schema for intratumoral flow cytometry analysis. A. Doublets and dead cells were removed from analysis, and unstained controls were consistently used to set analysis gates (bottom left panel). B. Gating structure for CD45^pos^ leukocyte analysis (left), which was used to set fluorescence minus one (FMO) control for CD19 staining (right). C. CD19^pos^ staining after initially gating on CD45^pos^ leukocytes.

**Supplemental Figure 2.**
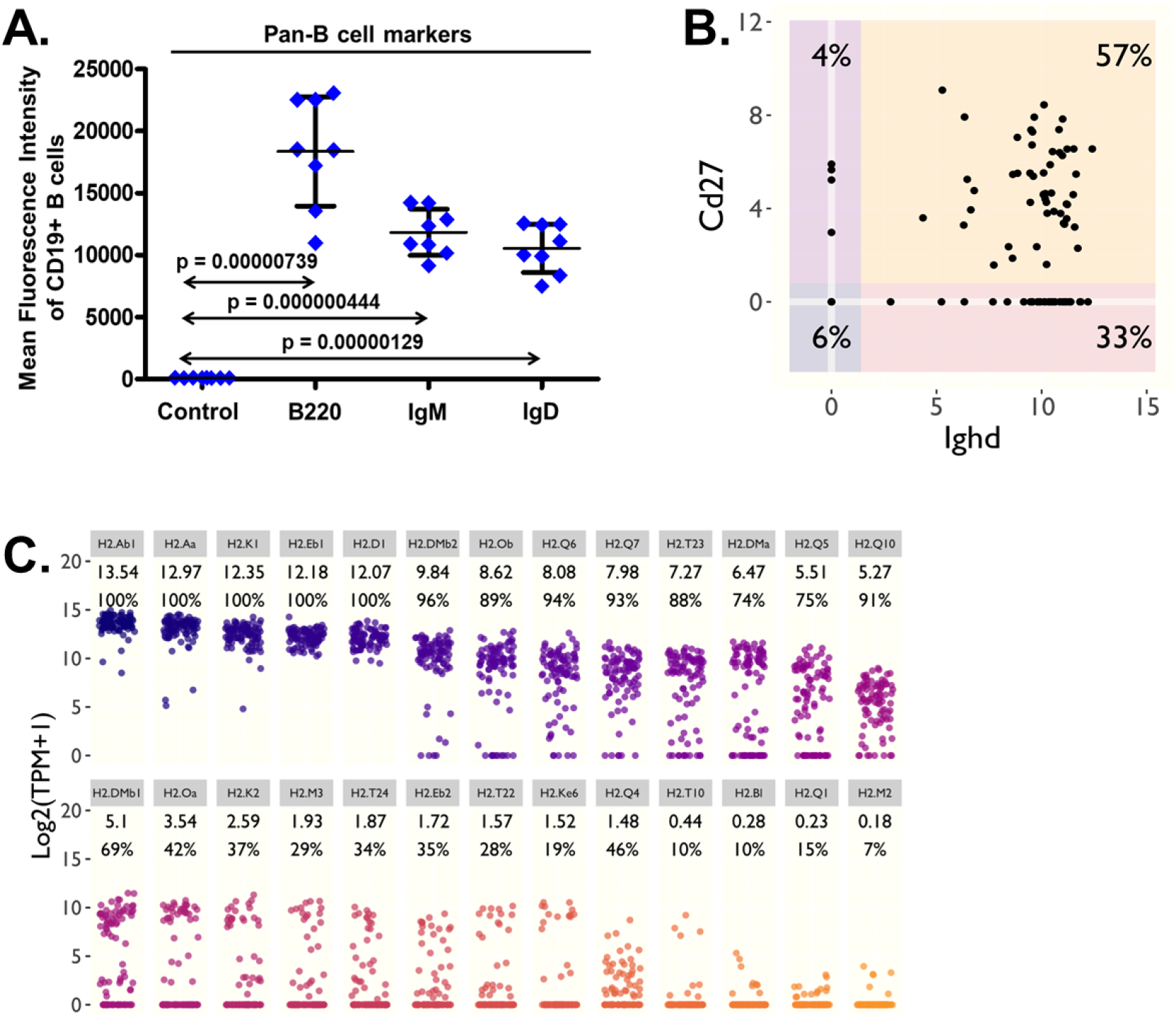
A. CD45^pos^CD19^pos^ TIL-B cells from B16F10 bearing mice were analyzed on day 13 for B cell markers. B-C. TIL-B cells were analyzed by scRNAseq data as in Fig. 2, except for (B.) *Ighd* and *Cd27* (C.) the presence of MHC genes.

**Supplemental Figure 3.**
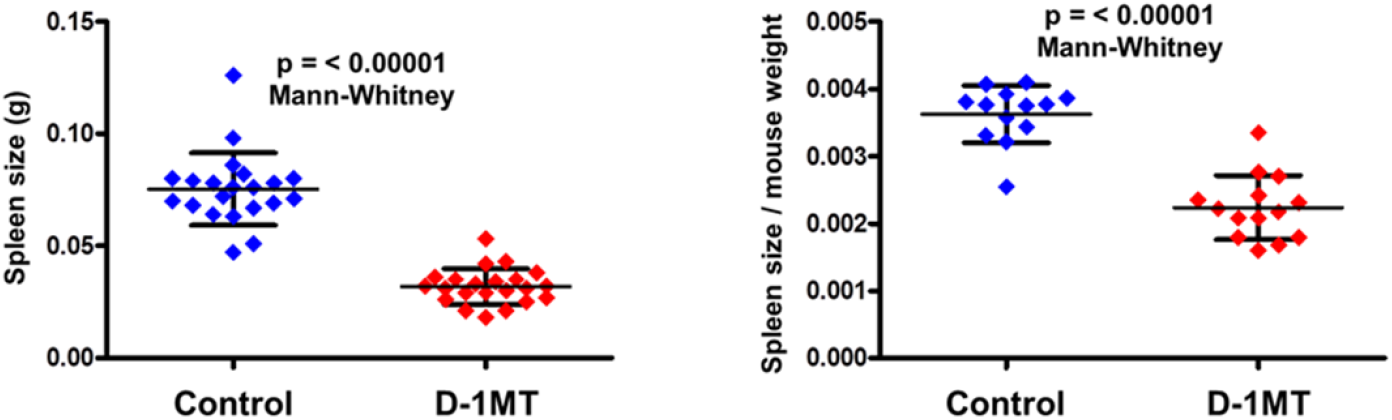
Melanoma bearing mice were treated as in **Fig. 1A**. On day 13 spleens were weighed (left) and normalized to total mouse weight (right).

**Supplemental Figure 4.**
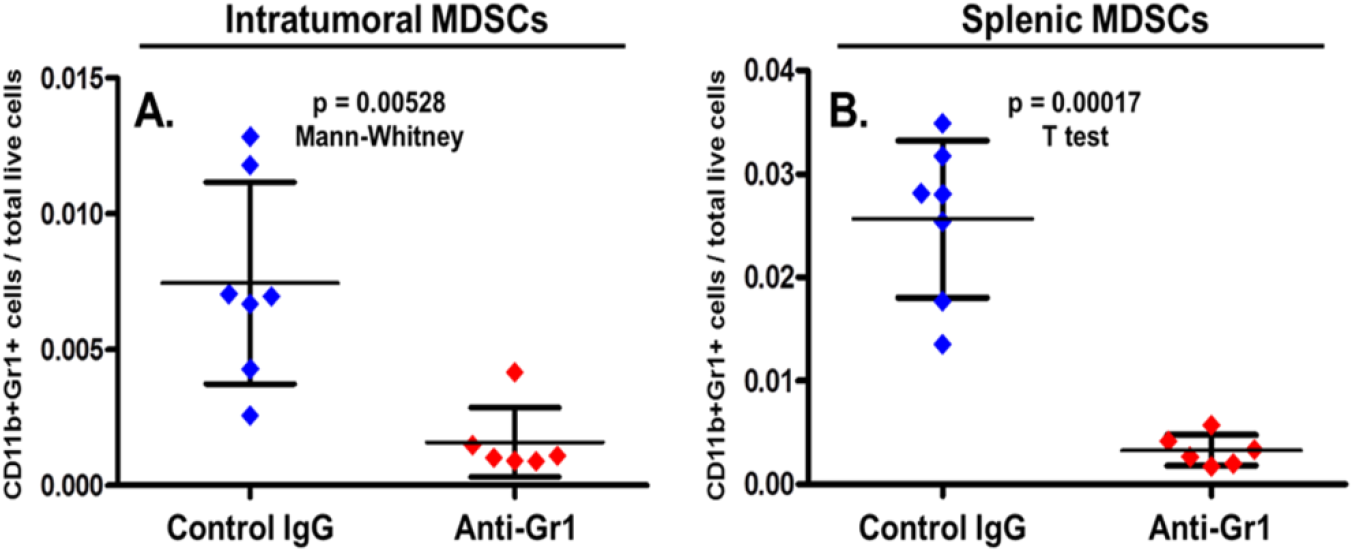
Melanoma-bearing mice were treated as in **Fig. 6C**. On day 13 tumors (A.) and spleens (B.) were analyzed for CD11b^pos^Gr1^pos^ MDSCs.

**Supplemental Figure 5.**
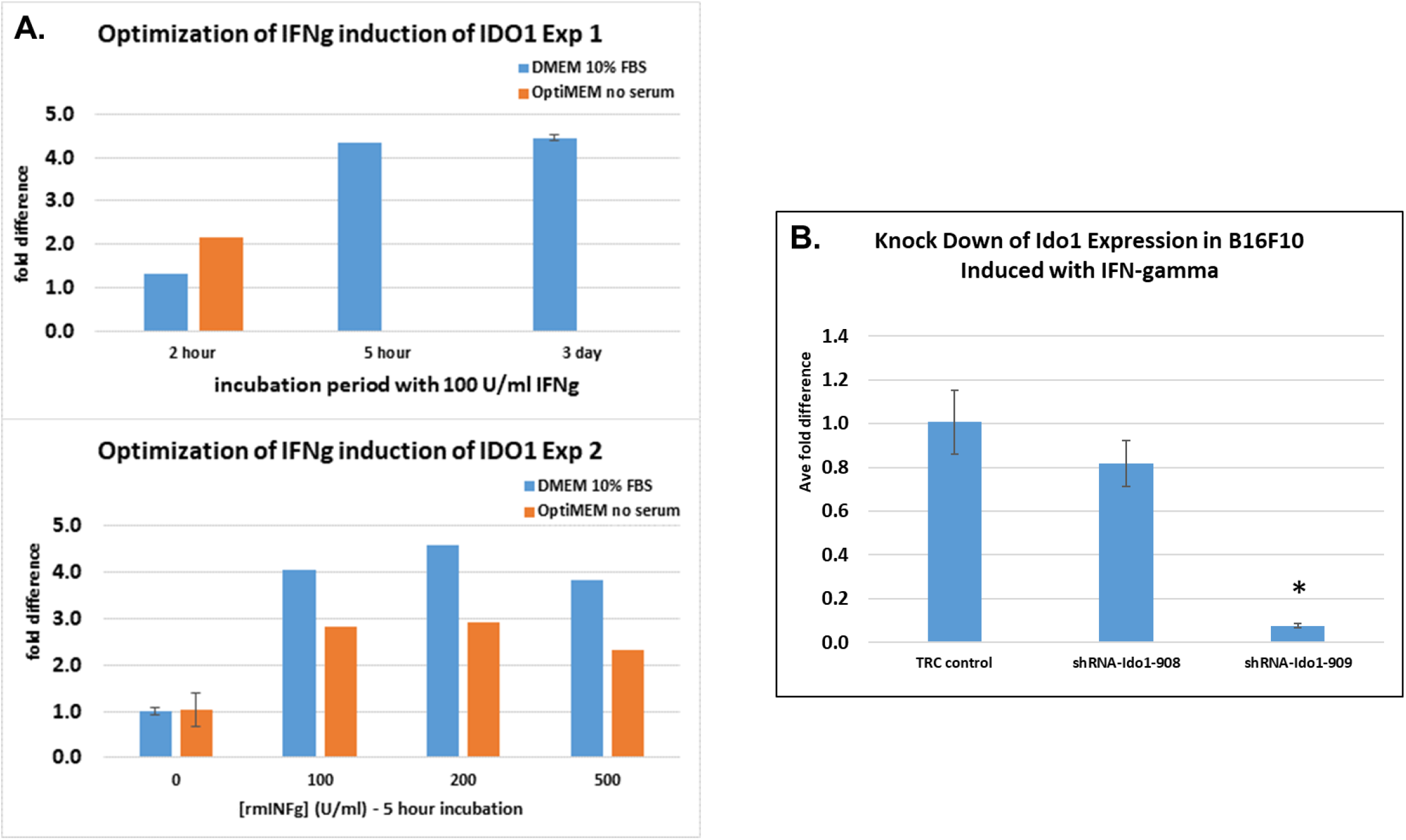
A, top. B16F10 melanoma cells were treated in vitro with 100 units/mL interferon gamma for 5 hours or 3 days then harvested and analyzed for *Ido1* expression by PCR. A, bottom. B16F10 melanoma cells were incubated with 100, 200, or 500 units/mL interferon gamma in media with or without serum as shown. After 5 hours tumor cells were harvested and analyzed for *Ido1* expression by PCR. B. *Ido1* expression levels of vector control and two different *Ido1* knockdown cells following induction with interferon gamma for 5 hours. The 909 *Ido1* knockdown was used for all *in vivo* experiments.

